# Intergenerational adaptations to stress are evolutionarily conserved, stress-specific, and have deleterious trade-offs

**DOI:** 10.1101/2021.05.07.443118

**Authors:** Nicholas O. Burton, Alexandra Willis, Kinsey Fisher, Fabian Braukmann, Jon Price, Lewis Stevens, L. Ryan Baugh, Aaron Reinke, Eric A. Miska

## Abstract

Despite reports of parental exposure to stress promoting physiological adaptations in progeny in diverse organisms, there remains considerable debate over the significance and evolutionary conservation of such multigenerational effects. Here, we investigate four independent models of intergenerational adaptations to stress in *C. elegans* – bacterial infection, eukaryotic infection, osmotic stress and nutrient stress – across multiple species. We found that all four intergenerational physiological adaptations are conserved in at least one other species, that they are stress-specific, and that they have deleterious trade-offs in mismatched environments. By profiling the effects of parental bacterial infection and osmotic stress exposure on progeny gene expression across species we established a core set of 279 highly conserved genes that exhibited intergenerational changes in expression in response to stress in all species tested and provide evidence suggesting that presumed adaptive and deleterious intergenerational effects are molecularly related at the gene expression level. By contrast, we found that these same stresses did not elicit any similarly conserved transgenerational changes in progeny gene expression three generations after stress exposure. We conclude that intergenerational responses to stress play a substantial and evolutionarily conserved role in regulating animal physiology and that the vast majority of the effects of parental stress on progeny gene expression are reversible and not maintained transgenerationally.

## Introduction

Multigenerational effects of a parent’s environment on progeny have been reported to contribute to numerous organismal phenotypes and pathologies in species ranging from plants to mammals (Agrawal et al., 1999; Bozler et al., 2019; Burton et al., 2020, 2017; Dantzer et al., 2013; Dias and Ressler, 2014; Hibshman et al., 2016; Houri-Zeevi et al., 2020; Jordan et al., 2019; Kaletsky et al., 2020; Kishimoto et al., 2017; Klosin et al., 2017; Luna et al., 2012; Ma et al., 2019; Moore et al., 2019; Öst et al., 2014; Palominos et al., 2017; Posner et al., 2019; Veenendaal et al., 2013; Vellichirammal et al., 2017; Webster et al., 2018; Wibowo et al., 2016; Willis et al., 2021).

These effects on progeny include many notable observations of intergenerational (lasting 1-2 generations) adaptive changes in phenotypically plastic traits such as the development of wings in pea aphids (Vellichirammal et al., 2017), helmet formation in *Daphnia* (Agrawal et al., 1999), accelerated growth rate in red squirrels (Dantzer et al., 2013), and physiological adaptations to osmotic stress and pathogen infection in both *Arabidopsis* (Luna et al., 2012; Wibowo et al., 2016) and *Caenorhabditis elegans* (Burton et al., 2020, 2017). These intergenerational adaptive changes in development and physiology can lead to substantial increases in organismal survival, with up to 50-fold increases in the survival of offspring from stressed parents being reported when compared to the offspring from naïve parents (Burton et al., 2020). While many of the most studied intergenerational effects of a parent’s environment on offspring have been identified in plants and invertebrates, intergenerational effects have also been reported in mammals (Dantzer et al., 2013; Dias and Ressler, 2014). Similar to findings in plants and invertebrates, some observations of intergenerational effects in mammals have been found to be physiologically adaptive (Dantzer et al., 2013), but many others, such as observations of fetal programming in humans (de Gusmão Correia et al., 2012; Langley-Evans, 2006; Schulz, 2010) and studies of the Dutch Hunger Winter (Veenendaal et al., 2013), have been reported to be deleterious. Nonetheless, even for these presumed deleterious intergenerational effects it has been hypothesized that under different conditions the intergenerational effects of fetal programming, such as the effects caused by the Dutch Hunger Winter, might be considered physiologically adaptive (Hales and Barker, 2001, 1992).

If intergenerational responses to environmental stresses represent evolutionarily conserved processes, if they are general or stress-specific effects, and whether adaptive and deleterious intergenerational effects are molecularly related remains unknown. Furthermore, multiple different studies have recently reported that some environmental stresses elicit changes in progeny physiology and gene expression that persist for three or more generations, also known as transgenerational effects (Kaletsky et al., 2020; Klosin et al., 2017; Ma et al., 2019; Moore et al., 2019; Posner et al., 2019; Webster et al., 2018). However, if intergenerational effects (lasting 1-2 generations) and transgenerational effects (lasting 3+ generations) represent related or largely separable phenomena remains unclear. Answering these questions is critically important not only in understanding the role that multigenerational effects play in evolution, but also in understanding how such effects might contribute to multiple human pathologies that have been linked to the effects of a parent’s environment on offspring, such as Type 2 diabetes and cardiovascular disease (Langley-Evans, 2006).

Here, we investigated the evolutionary conservation, stress specificity, and potential tradeoffs of four independent models of intergenerational adaptations to stress in *C. elegans –* bacterial infection, eukaryotic infection, nutrient stress and osmotic stress. We found that all four models of intergenerational adaptive effects are conserved in at least one other species, but that all exhibited a different pattern of evolutionary conservation. Each intergenerational adaptive effect was stress-specific and multiple intergenerational adaptive effects exhibited deleterious tradeoffs in mismatched environments or environments where multiple stresses were present simultaneously. By profiling the effects of multiple different stresses on offspring gene expression across species we identified a set of 279 genes that exhibited intergenerational changes in gene expression in response to stress in all species tested. In addition, we found that an inversion in the expression of a subset of these genes, from increased expression to decreased expression in the offspring of stressed parents, correlates with an inversion of an adaptive intergenerational response to bacterial infection in *C. elegans* and *C. kamaaina* to a deleterious intergenerational effect in *C. briggsae*. Lastly, we report that the vast majority of the intergenerational effects of multiple different stresses on offspring gene expression were not maintained transgenerationally in F3 progeny and that no transgenerational changes in gene expression that were observed in *C. elegans* were conserved in in a second *Caenorhabditis* species that exhibits phenotypically conserved intergenerational responses to stress (*C. kamaaina*). Our findings demonstrate that intergenerational adaptive responses to stress are evolutionarily conserved, stress-specific, and likely represent a distinct phenomenon from transgenerational effects. In addition, our findings suggest that the mechanisms that mediate intergenerational adaptive responses in some species might be related to the mechanisms that mediate intergenerational deleterious effects in other species.

## Results

### Intergenerational adaptations to stress are evolutionarily conserved

To test if any of the intergenerational adaptations to stress that have been reported in *C. elegans* are evolutionarily conserved in other species we focused on four recently described intergenerational adaptations to abiotic and biotic stresses - osmotic stress (Burton et al., 2017), nutrient stress (Hibshman et al., 2016; Jordan et al., 2019), *Pseudomonas vranonvensis* infection (bacterial) (Burton et al., 2020), and *Nematocida parisii* infection (eukaryotic – microsporidia) (Willis et al., 2021). We tested if these four intergenerational adaptive responses were conserved in four different species of *Caenorhabditis* (*C. briggsae, C. elegans, C. kamaaina,* and *C. tropicalis*) that shared a last common ancestor approximately 30 million years ago (Figure 1A) (Cutter, 2008). These species were chosen because they represent multiple independent branches of the *Elegans* group (Figure 1A) and because we could probe the conservation of underlying mechanisms using established genetics approaches.

**Figure 1.**
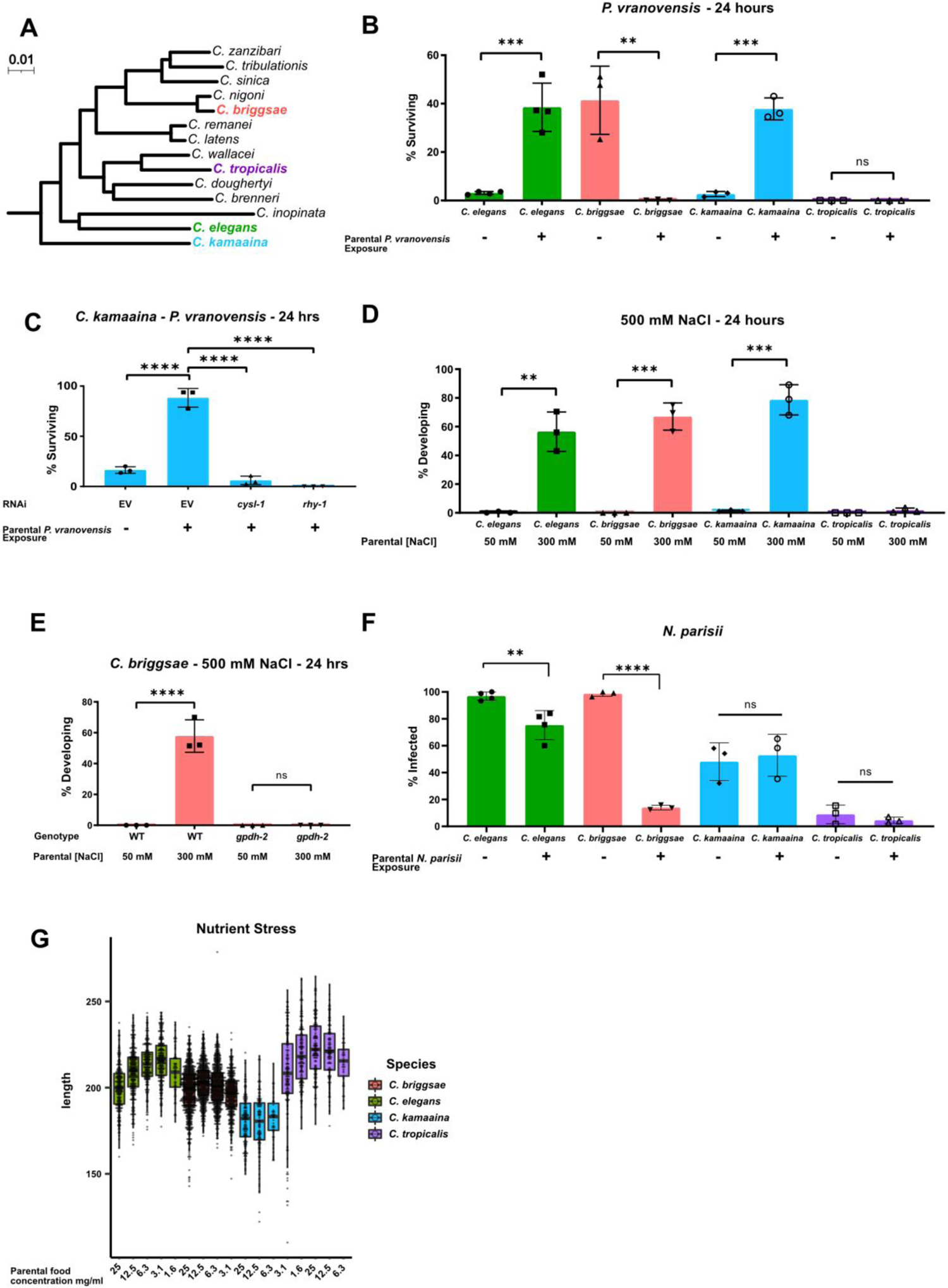
Intergenerational adaptations to multiple stresses are evolutionarily conserved in multiple species of *Caenorhabditis*. (A) Phylogenetic tree of the *Elegans* group of *Caenorhabditis* species adapted from Stevens et al., 2020. Scale represents substitutions per site. (B) Percent of wild-type *C. elegans* (N2), *C. kamaaina* (QG122), *C. briggsae* (AF16), and *C. tropicalis* (JU1373) animals surviving after 24 hours on plates seeded with *P. vranovensis* BIGb0446. Data presented as mean values ± s.d. *n* = 3–4 experiments of >100 animals. (C) Percent of *C. kamaaina* wild-type (QG122) animals surviving after 24 hours of exposure to *P. vranovensis*. Data presented as mean values ± s.d. *n* = 3 experiments of >100 animals. (D) Percent of wild-type animals mobile and developing at 500 mM NaCl after 24 hours. Data presented as mean values ± s.d. *n* = 3 experiments of >100 animals. (E) Percent of wild-type and *Cbr-gpdh-2(syb2973)* mutant *C. briggsae* (AF16) mobile and developing after 24 hours at 500 mM NaCl. Data presented as mean values ± s.d. *n* = 3 experiments of >100 animals. (F) Percent of animals exhibiting detectable infection by *N. parisii* as determined by DY96 staining after 72 h for *C. elegans* and *C. briggsae*, or 96 h for *C. kamaaina* and *C. tropicalis.* Data presented as mean values ± s.e.m. *n* = 3-4 experiments of 83-202 animals. (G) Boxplots for length of L1 progeny from P0 parents that were subject to the HB101 dose series. Larvae were measured using Wormsizer. Boxplots show median length with four quartiles. n = 3-8 experiments of 50-200 animals. \*\**p* < 0.01, *** *p* < 0.0001, \*\*\*\**p* < 0.0001

We exposed parents of all four species to *P. vranovensis* and subsequently studied their offspring’s survival rate in response to future *P. vranovensis* exposure. We found that parental exposure to the bacterial pathogen *P. vranovensis* protected offspring from future infection in both *C. elegans* and *C. kamaaina* (Figure 1B) and that this adaptive intergenerational effect in *C. kamaaina* required the same stress response genes (*cysl-1* and *rhy-1*) as previously reported for *C. elegans* (Burton et al., 2020) (Figure 1C), indicating that these animals intergenerationally adapt to infection via a similar and potentially conserved mechanism. By contrast, we found that naïve *C. briggsae* animals were more resistant to *P. vranovensis* than any of the other species tested, but exposure of *C. briggsae* parents to *P. vranovensis* caused greater than 99% of offspring to die upon future exposure to *P. vranovensis* (Figure 1B). We confirmed that parental *P. vranovensis* exposure resulted in an adaptive intergenerational effect for *C. elegans* but a deleterious intergenerational effect for *C. briggsae* by testing multiple additional wild isolates of both species (Supplemental Figure 1A-C). Parental exposure to *P. vranovensis* had no observable effect on offspring response to infection in *C. tropicalis* (Figure 1B). We conclude that parental exposure to *P. vranovensis* causes substantial changes in offspring susceptibility to future *P. vranovensis* exposure in multiple species, but whether those effects are protective or deleterious for offspring is species dependent.

Using a similar approach to investigate intergenerational adaptive responses to other stresses, we found that parental exposure to mild osmotic stress protected offspring from future osmotic stress in all of *C. elegans, C. briggsae*, and *C. kamaaina,* but again not in *C. tropicalis* (Figure 1D). This intergenerational adaptation to osmotic stress in *C. briggsae* and *C. kamaaina* required the glycerol-3-phosphate dehydrogenase *gpdh-2* (Figure 1E and Supplemental Figure 1D), similar to previous observations for *C. elegans* (Burton et al., 2017) and indicating that these adaptations are regulated by similar and likely evolutionarily conserved mechanisms.

We then sought to test if intergenerational resistance to infection by the eukaryotic pathogen *N. parisii* is similarly conserved in *Caenorhabditis* species. *N. parisii* is a common natural pathogen of both *C. elegans* and *C. briggsae* (Zhang et al., 2016). Here, we show that *N. parisii* can also infect *C. kamaaina* and *C. tropicalis* (Supplemental Figure 2). By investigating the effects of parental *N. parisii* infection on offspring across species, we found that parental exposure of *C. elegans* and *C. briggsae* to *N. parisii* protected offspring from future infection (Figure 1F). By contrast, parental exposure of *C. kamaaina* and *C. tropicalis* to *N. parisii* had no observable effect on offspring infection rate (Figure 1F).

Lastly, we investigated the intergenerational effects of nutrient stress on offspring. We found that parental nutrient stress by food deprivation resulted in larger offspring in both *C. elegans* and *C. tropicalis*, which is predicted to be adaptive (Hibshman et al., 2016), but had minimal effects on offspring size in *C. briggsae* and *C. kamaaina* (Figure. 1G). Collectively, our findings indicate that all four reported intergenerational adaptive effects in *C. elegans* are conserved in at least one other species but all four show a different pattern of conservation, which is consistent with each response being regulated by distinct molecular mechanisms (Burton et al., 2020, 2017; Hibshman et al., 2016; Jordan et al., 2019; Willis et al., 2021).

### The effects of parental bacterial infection on offspring gene expression correlate with offspring pathogen sensitivity

Of the four intergenerational models investigated here, parental exposure of *C. elegans* to osmotic stress and *P. vranovensis* infection were previously reported to cause substantial changes in offspring gene expression, including the expression of genes that are required for the observed intergenerational adaptations (Burton et al., 2020, 2017). We exposed *C. elegans, C. briggsae, C. kamaaina,* and *C. tropicalis* to osmotic stress and *P. vranovensis* infection and subsequently performed RNA-seq on offspring to test: (1) if the specific heritable changes in gene expression in response to each of these stresses are conserved across species and (2) if any changes in gene expression correlate with the phenotypic differences in intergenerational responses to stress we observed in the different species. This analysis allowed us to compare the effects of parental stress on offspring gene expression of 7,587 single-copy orthologues that are conserved across all four species (Supplemental Table 1).

Consistent with previous observations in *C. elegans*, we found that parental exposure to *P. vranovensis* resulted in substantial changes in offspring gene expression in all four species we investigated (Figure 2A-D and Supplemental Table 2). Of the 7,587 single copy orthologues shared between the four species, we found 2,663 genes that exhibited differential expression in the offspring of infected animals in *C. elegans* and at least one other species (Figure 2D and Supplemental Table 2). Furthermore, we found that 275 genes are differentially expressed in the offspring of parents exposed to *P. vranovensis* in all four species (Figure 2D and Supplemental Table 2). These data indicate that parental exposure to the bacterial pathogen *P. vranovensis* leads to changes in offspring gene expression at a common set of stress response genes in diverse species of *Caenorhabditis*.

**Figure 2.**
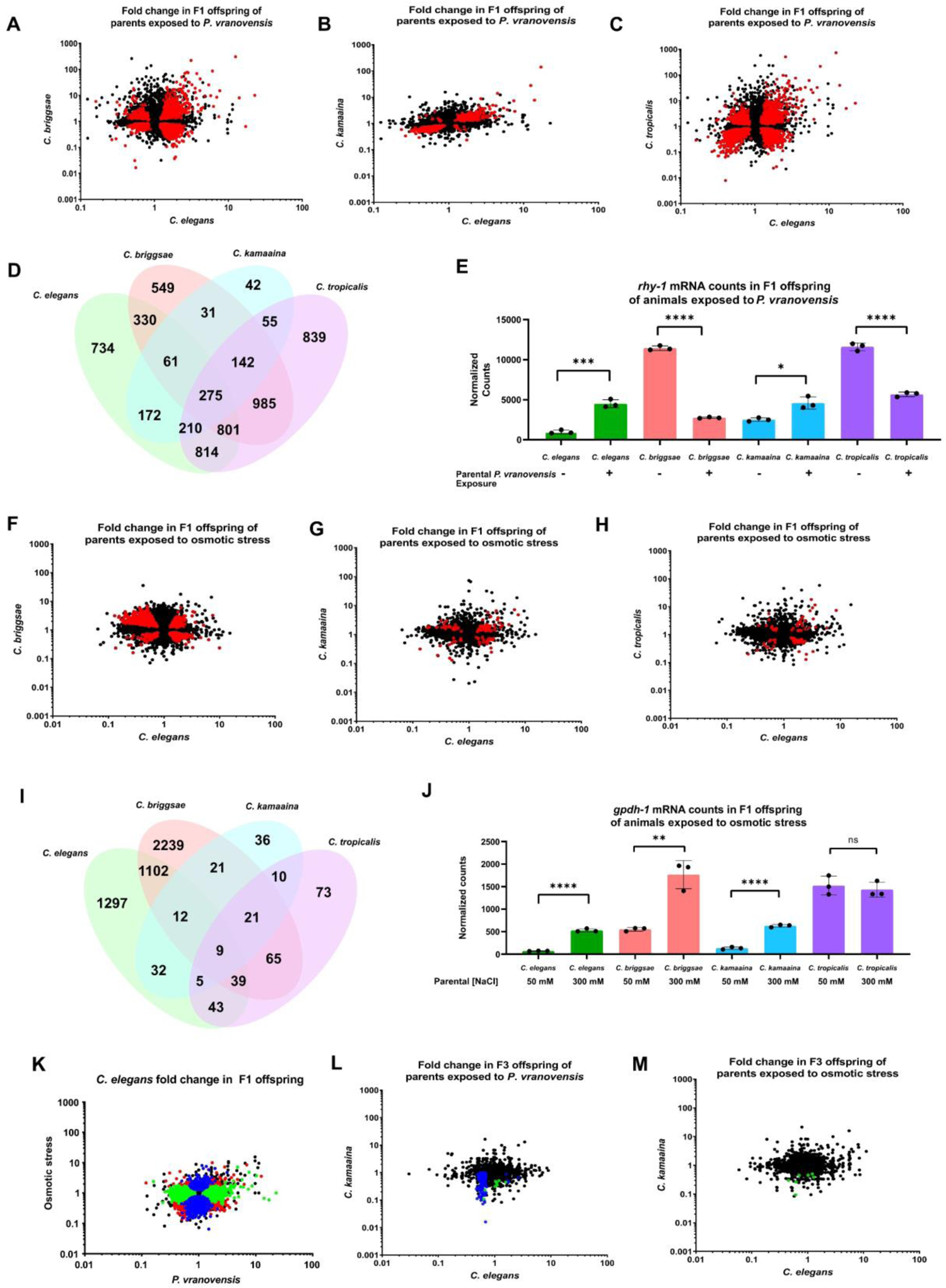
Parental exposure to *P. vranovensis* and osmotic stress have overlapping effects on offspring gene expression across multiple species. (A) Average fold change of 7,587 single-copy orthologue genes in F1 progeny of *C. elegans* and *C. briggsae* parents fed *P. vranovensis* BIGb0446 when compared to parents fed *E. coli* HB101. Average fold change from three replicates. Red dots represent genes that exhibit statistically significant (*padj* < 0.01) changes in both species. (B) Average fold change of 7,587 single-copy orthologue genes in F1 progeny of *C. elegans* and *C. kamaaina* parents fed *P. vranovensis* BIGb0446 when compared to parents fed *E. coli* HB101. Average fold change from three replicates. Red dots represent genes that exhibit statistically significant (*padj* < 0.01) changes in both species (C) Average fold change of 7,587 single-copy orthologue genes in F1 progeny of *C. elegans* and *C. tropicalis* parents fed *P. vranovensis* BIGb0446 when compared to parents fed *E. coli* HB101. Average fold change from three replicates. Red dots represent genes that exhibit statistically significant (*padj* < 0.01) changes in both species (D) Venn diagram of the number of genes that exhibit overlapping statistically significant (*padj* < 0.01) changes in expression in F1 progeny of animals exposed to *P. vranovensis* BIGb0446 in each species. (E) Normalized counts of reads matching orthlougues of *rhy-1* in the F1 offspring of parents fed either *E. coli* HB101 or *P. vranovensis* BIGb0446. Data from Supplemental Table 2. *n* = 3 replicates. (F) Average fold change of 7,587 single-copy orthologue genes in F1 progeny of *C. elegans* and *C. briggsae* parents grown at 300 mM NaCl when compared to parents grown at 50 mM NaCl. Average fold change from three replicates. Red dots represent genes that exhibit statistically significant (*padj* < 0.01) changes in both species. (G) Average fold change of 7,587 single-copy orthologue genes in F1 progeny of *C. elegans* and *C. kamaaina* parents grown at 300 mM NaCl when compared to parents grown at 50 mM NaCl. Average fold change from three replicates. Red dots represent genes that exhibit statistically significant (*padj* < 0.01) changes in both species. (H) Average fold change of 7,587 single-copy orthologue genes in F1 progeny of *C. elegans* and *C. tropicalis* parents grown at 300 mM NaCl when compared to parents grown at 50 mM NaCl. Average fold change from three replicates. Red dots represent genes that exhibit statistically significant (*padj* < 0.01) changes in both species. (I) Venn diagram of the number of genes that exhibit overlapping statistically significant (*padj* < 0.01) changes in expression in F1 progeny of animals grown at 300 mM NaCl in each species. (J) Normalized counts of reads matching orthlougues of *gpdh-1* in the F1 progeny of parents grown at either 300 mM NaCl or 50 mM NaCl. Data from Supplemental Table 3. *n* = 3 replicates. (K) Average fold change for 7,587 orthologue genes in F1 progeny of *C. elegans* parents fed *P. vranovensis* or exposed to 300 mM NaCl when compared to naïve parents. Average fold change from three replicates. Red dots – genes that change in expression in response to both stresses. Blue dots – genes that change in expression in response to only osmotic stress. Green dots – genes that change in expression in response to only *P. vranovensis.* (L) Average fold change of 7,512 single-copy orthologue genes in F3 progeny of *C. elegans* and *C. kamaaina* fed *P. vranovensis* BIGb0446 when compared to those fed *E. coli* HB101. Average fold change from three replicates. Blue dots represent genes that exhibited statistically significant (*padj* < 0.01) changes in *C. elegans*. Green dots represent genes that exhibited statistically significant (*padj* < 0.01) changes in *C. kamaaina.* (M) Average fold change of 7,512 single-copy orthologue genes in F1 progeny of *C. elegans* and *C. kamaaina* parents grown at 300 mM NaCl when compared to parents grown at 50 mM NaCl. Average fold change from three replicates. Green dots represent genes that exhibited statistically significant (*padj* < 0.01) changes in *C. kamaaina. *p<*0.05, \*\**p* < 0.01, *** *p* < 0.0001, \*\*\*\**p* < 0.0001

Parental exposure of *C. elegans* and *C. kamaaina* to *P. vranovensis* led to increased progeny resistance to future *P. vranovensis* exposure (Figure 1B). By contrast, parental exposure of *C. briggsae* to *P. vranovensis* led to increased offspring susceptibility to *P. vranovensis* (Figure1B). We hypothesized that differences in the expression of genes previously reported to be required for adaptation to *P. vranovensis*, such as the acyltransferase *rhy-1*, might underlie these differences between species. We therefore investigated whether any genes exhibited specific changes in expression in *C. elegans* and *C. kamaaina* that were either absent or inverted in *C. briggsae.* We found that of the 3,397 genes that exhibited differential expression in the offspring of parents exposed to *P. vranovensis* in *C. elegans*, only 718 were also differentially expressed in *C. kamaaina* (Supplemental Table 2). From this refined list of 718 genes, we found that 287 exhibited increased expression in both *C. elegans* and *C. kamaaina*. Of these 287 genes, 66 were not differentially expressed in *C. briggsae* and 52 exhibited decreased expression in the offspring of *C. briggsae* parents exposed to *P. vranovensis* (Supplemental Table 2). Similarly, we identified 405 genes that exhibited decreased expression in both the offspring of *C. elegans* and *C. kamaaina* parents exposed to *P. vranovensis*. Of these genes, 303 were not differentially expressed in *C. briggsae* and 18 exhibited increased expression in the offspring of *C. briggsae* parents exposed to *P. vranovensis* (Supplemental Table 2). These results indicate that a majority of the genes that are differentially expressed in the offspring of both *C. elegans* and *C. kamaaina* either do not change in *C. briggsae* or change in the opposite direction.

Three genes, *cysl-1, cysl-2* and *rhy-1*, were previously reported to be required for *C. elegans* to intergenerationally adapt to *P. vranovensis* (Burton et al., 2020). Here, we found that all three genes exhibit significantly increased expression in the offspring of infected parents in both *C. elegans* and *C. kamaaina*. By contrast, *rhy-1* exhibited a 4-fold decrease in expression in *C. briggsae* offspring from infected parents (Figure 2E). Similarly, we found that parental exposure of *C. briggsae* to *P. vranovensis* had either no effect or a substantially reduced effect on the expression of *cysl-1* and *cysl-2* in the offspring of infected parents when compared to *C. elegans* and *C. kamaaina* (Supplemental Table 2). Notably, the directional change of *rhy-1* expression in progeny of animals exposed to *P. vranovensis* correlates with the observation that parental exposure to *P. vranovensis* results in enhanced pathogen resistance in offspring in *C. elegans* and *C. kamaaina* but has a strong deleterious effect on pathogen resistance in *C. briggsae* (Figure 1B). These findings suggest that molecular mechanisms underlying adaptive and deleterious effects in different species might be related and dependent on the direction of changes in gene expression of specific stress response genes.

### Different parental stresses have distinct effects on offspring gene expression

We performed the same analysis on the offspring of all four species from parents exposed to osmotic stress. From this analysis we observed that parental exposure to osmotic stress resulted 1,163 genes exhibiting differential expression in both *C. elegans* and *C. briggsae* offspring (Figure 2F-K and Supplemental Table 3). In addition, we found that these changes in gene expression were largely distinct from the gene expression changes observed in the offspring of parents exposed to *P. vranovensis* (Figure 2K and Supplemental Tables 2 and 3), indicating that different parental stresses have distinct effects on offspring gene expression. However, parental exposure to *C. kamaaina* and *C. tropicalis* to osmotic stress resulted in approximately 5-fold fewer changes in offspring gene expression (Figure 2G-H and Supplemental Table 3). In total only nine genes exhibited differential expression in the offspring of parents exposed to osmotic stress in all four species (Figure 2I) and five of these nine were also observed to change in the offspring of parents exposed to *P. vranovensis* (Supplemental Table 3).

Unlike *C. elegans, C. briggsae*, and *C. kamaaina*, parental exposure of *C. tropicalis* to osmotic stress did not protect offspring from future osmotic stress (Figure 1D). We therefore identified genes that were differentially expressed in the F1 offspring of *C. elegans*, *C. briggsae,* and *C. kamaaina* exposed to osmotic stress, but not in *C. tropicalis.* From this analysis we identified eleven genes that are specifically differentially expressed in the three species that adapt to osmotic stress but not in *C. tropicalis*; this list of genes includes the glycerol-3-phosphate dehydrogenase *gpdh-1* which is one of the most upregulated genes in response to osmotic stress and is known to be required for animals to properly respond to osmotic stress (Lamitina et al., 2006) (Figure 2J). These results suggest that, similar to our observations for *P. vranovensis* infection, different patterns in the expression of osmotic stress response genes correlate with different intergenerational phenotypic responses to osmotic stress.

Differences in gene expression in the offspring of stressed parents could be due to programmed changes in expression in response to stress or due to indirect effects caused by changes in developmental timing. To confirm that the embryos from all conditions were collected at the same developmental stage we compared our RNA-seq findings to a time resolved transcriptome of *C. elegans* development (Boeck et al., 2016). Consistent with our visual observations that a vast majority of offspring collected were in the comma stage of embryo development, we found that the gene expression profiles of all offspring from both naïve and stressed parents overlapped strongly with the 330-450 minute timepoints of development (Supplemental Figure 3A). In addition, we found that approximately 50% of all genes that were differentially expressed in the offspring of stressed parents when compared to naïve parents exhibited a change in gene expression that was more than one standard deviation outside their average expression across all timepoints of embryo development (Supplemental Figure 3B-3C). We similarly found that many of the genes known to be required for intergenerational responses to stress exhibit expression that is outside the range of expression observed at any time point of early development (Supplemental Figure 3D-3E). These results suggest that a majority of the expression differences we observed in the offspring of stressed parents were not due to differences in developmental timing.

**Figure 3.**
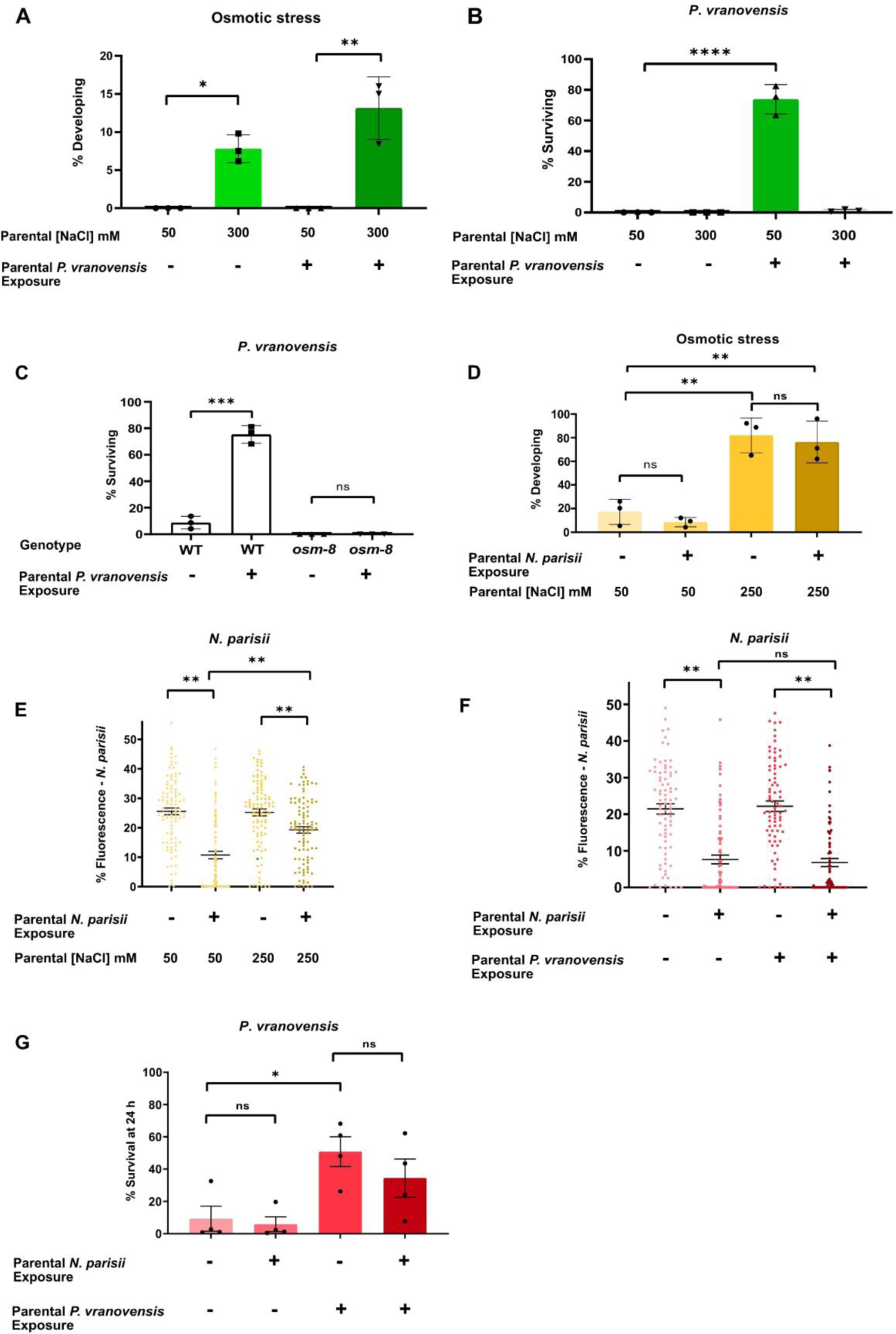
Intergenerational adaptations to stress are stress-specific and have deleterious trade-offs. (A) Percent of wild-type *C. elegans* mobile and developing at 500 mM NaCl after 24 hours. Data presented as mean values ± s.d. *n* = 3 experiments of >100 animals. (B) Percent of wild-type *C. elegans* surviving after 24 hours of exposure to *P. vranovensis* BIGb0446. Data presented as mean values ± s.d. *n* = 3 experiments of >100 animals. (C) Percent of wild-type and *osm-8(n1518) C. elegans* surviving after 24 hours of exposure to *P. vranovensis* BIGb0446. Data presented as mean values ± s.d. *n* = 3 experiments of >100 animals. (D) Percent of wild-type *C. elegans* mobile and developing at 420 mM NaCl after 48 hours. Data presented as mean values ± s.d. *n* = 3 experiments of >100 animals. (E) *N. parisii* parasite burden of individual *C. elegans* after 72 h (as determined by percentage fluorescence from DY96-stained spores after 72 h). Data presented as mean values ± s.e.m. *n =* 4 experiments of 25 animals (F) *N. parisii* parasite burden of individual *C. elegans* after 72 h (as determined by percentage fluorescence from DY96-stained spores after 72 h). Data presented as mean values ± s.e.m. *n =* 3 experiments of 25 animals. (G) Percent of wild-type *C. elegans* surviving after 24 hours of exposure to *P. vranovensis* BIGb0446. Data presented as mean values ± s.e.m. *n* = 3 experiments of >100 animals. *\*p<*0.05, \*\**p* < 0.01, *** *p* < 0.0001, \*\*\*\**p* < 0.0001

### The effects of parental bacterial infection and osmotic stress on offspring gene expression are not maintained transgenerationally

Determining whether the effects of parental exposure to stress on offspring gene expression are reversible after one generation or if any changes in gene expression persist transgenerationally is a critical and largely unanswered question in the field of multigenerational effects. To test if any of the intergenerational changes in gene expression that we observed persist transgenerationally we performed RNA-seq of F3 progeny of *C. elegans* exposed to both *P. vranovensis* and osmotic stress. We found that only 121 of the 3,397 genes that exhibited intergenerational (F1) changes in gene expression in response to *P. vranovensis* infection were also differentially expressed transgenerationally in *C. elegans* F3 progeny (Figure 2K and Supplemental Table 4).

Furthermore, we found that only two of the 2,539 genes that exhibited intergenerational changes in expression in response to osmotic stress were also differentially expressed in *C. elegans* F3 progeny (Figure 2L and Supplemental Table 4). We conclude that the vast majority of intergenerational effects of these stresses on gene expression do not persist transgenerationally. To test if any of the 123 genes that exhibited differential expression in the F3 progeny of *C. elegans* exposed to *P. vranovensis* or osmotic stress also exhibit transgenerational changes in expression in other *Caenorhabditis* species, we performed the same experiments on *C. kamaaina*, which also intergenerationally adapts to both *P. vranovensis* infection and osmotic stress. We found that none of the 123 genes that were differentially expressed in the F3 progeny of *C. elegans* exposed to either *P. vranovensis* or osmotic stress were also differentially expressed in the F3 progeny of *C. kamaaina* under the same conditions (Figure 2K and Supplemental Table 4). Our results suggest that neither of these biotic or abiotic stresses that elicit robust intergenerational changes in gene expression cause similar transgenerational changes in gene expression in a second *Caenorhabditis* species under the same conditions. We note, however, that it remains possible that transgenerational effects of these stresses could persist through other mechanisms, could affect the expression of genes that are not clearly conserved between species, or could exert weaker effects on broad classes of genes that would not be detectable at any specific individual loci as was reported for the transgenerational effects of starvation and loss of COMPASS complex function on gene expression in *C. elegans* (Greer et al., 2011; Webster et al., 2018). Furthermore, it is possible that transgenerational effects on gene expression in *C. elegans* are restricted to germ cells (Buckley et al., 2012; Houri-Zeevi et al., 2020; Posner et al., 2019) and are not detectable in somatic tissue. Such effects that occur specifically in germ cells might not have been detectable in the early developmental stage assayed here.

### Intergenerational responses to stress can have deleterious trade offs

Intergenerational changes in animal physiology that protect offspring from future exposure to stress could be stress-specific or could converge on a broadly stress resistant state. If intergenerational adaptive effects are stress-specific, then it is expected that parental exposure to a given stress will protect offspring from that same stress but potentially come at the expense of fitness in mismatched environments. If intergenerational adaptations to stress converge on a generally more stress resistant state, then parental exposure to one stress might protect offspring against many different types of stress. To determine if the intergenerational effects we investigated here represent specific or general responses we assayed how parental *C. elegans* exposure to osmotic stress, *P. vranovensis* infection, and *N. parisii* infection, either alone or in combination, affected offspring responses to mismatched stresses. We found that parental exposure to *P. vranovensis* did not affect the ability of animals to intergenerationally adapt to osmotic stress (Figure 3A). By contrast, parental exposure to osmotic stress completely eliminated the ability of animals to intergenerationally adapt to *P. vranovensis* (Figure 3B). This effect is unlikely to be due to the effects of osmotic stress on *P. vranovensis* itself, as mutant animals that constitutively activate the osmotic stress response (*osm-8*) were also completely unable to adapt to *P. vranovensis* infection (Figure. 3C) (Rohlfing et al., 2011). We conclude that animals’ intergenerational responses to *P. vranovensis* and osmotic stress are stress-specific, consistent with our observation that parental exposure to these two stresses resulted in distinct changes in offspring gene expression (Figure 2K).

We performed a similar analysis comparing animals’ intergenerational response to osmotic stress and the eukaryotic pathogen *N. parisii*. We previously reported that L1 parental infection with *N. parisii* results in progeny that is more sensitive to osmotic stress (Willis et al., 2021). Here we found that L4 parental exposure of *C. elegans* to *N. parisii* had a small, but not significant effect on offspring response to osmotic stress (Figure 3D). However, similar to our observations for osmotic stress and bacterial infection, we found that parental exposure to both osmotic stress and *N. parisii* infection simultaneously resulted in offspring that were less protected against future *N. parisii* infection than when parents are exposed to *N. parisii* alone (Figure 3E). Collectively, these data further support the conclusion that intergenerational responses to infection and osmotic stress are stress-specific and suggest that intergenerational adaptations to osmotic stress might come at the expense of animals’ ability to properly respond to bacterial or eukaryotic infections when either is paired with osmotic stress.

To compare animals’ intergenerational responses to bacterial infection and eukaryotic infection we performed a similar comparative analysis. We found that parental exposure to *P. vranovnesis* had no observable effect on offspring response to *N. parisii* either alone or when both pathogens were present simultaneously (Figure 1F). Similarly, we found that parental exposure to *N. parisii* had no observable effect on offspring response to *P. vranovensis* either alone or when both pathogens were present at the same time (Figure 1G). We conclude that intergenerational adaptations to osmotic stress, *P. vranovensis* infection and *N. parisii* infection are largely stress-specific.

### Intergenerational responses to *Pseudomonas* pathogens are distinct from other bacterial pathogens

To further probe the specificity of intergenerational responses to stress we also sought to determine if the substantial changes in pathogen resistance and gene expression observed in *C. elegans* offspring from parents exposed to the bacterial pathogen *P. vranovensis* were specific to this pathogen or part of a general response to bacterial pathogens. To test this we first screened wild bacterial isolates from France (Samuel et al., 2016) and the United Kingdom (Supplemental Table 5) for those that are potential natural pathogens of *C. elegans* and that also intergenerationally affect *C. elegans* survival or growth rate. From this analysis we identified a new *Pseudomonas* isolate, *Pseudomonas sp.* 15C5, where parental exposure to *Pseudomonas sp.* 15C5 enhanced offspring growth rate in response to future exposure to *Pseudomonas sp.* 15C5 (Figure 4A). This intergenerational effect resembled *C. elegans* intergenerational adaptation to *P. vranovensis* and we found that parental exposure to either isolate of *Pseudomonas* protected offspring from future exposure to the other *Pseudomonas* isolate (Figure 4A-B). To test if *Pseudomonas sp. 15C5* was a new isolate of *P. vranovensis* or a distinct species of *Pseudomonas* we performed both 16S rRNA sequencing and sequenced the gene *rpoD* of *Pseudomonas sp.* 15C5. From this analysis we found that *Pseudomonas sp.* 15C5 is not an isolate of *P. vranovensis* and is most similar to *Pseudomonas putida* – 99.49% identical 16S rRNA and 98.86% identical *rpoD* by BLAST (Supplemental File 1). These results indicate that parental exposure to multiple different *Pseudomonas* species can protect offspring from future pathogen exposure. We note, however, that other pathogenic species of *Pseudomonas*, such as *P. aeruginosa*, did not cross protect against *P. vranovensis* (Burton et al., 2020), indicating that not all pathogenic species of *Pseudomonas* result in the same intergenerational changes in offspring pathogen resistance.

**Figure 4.**
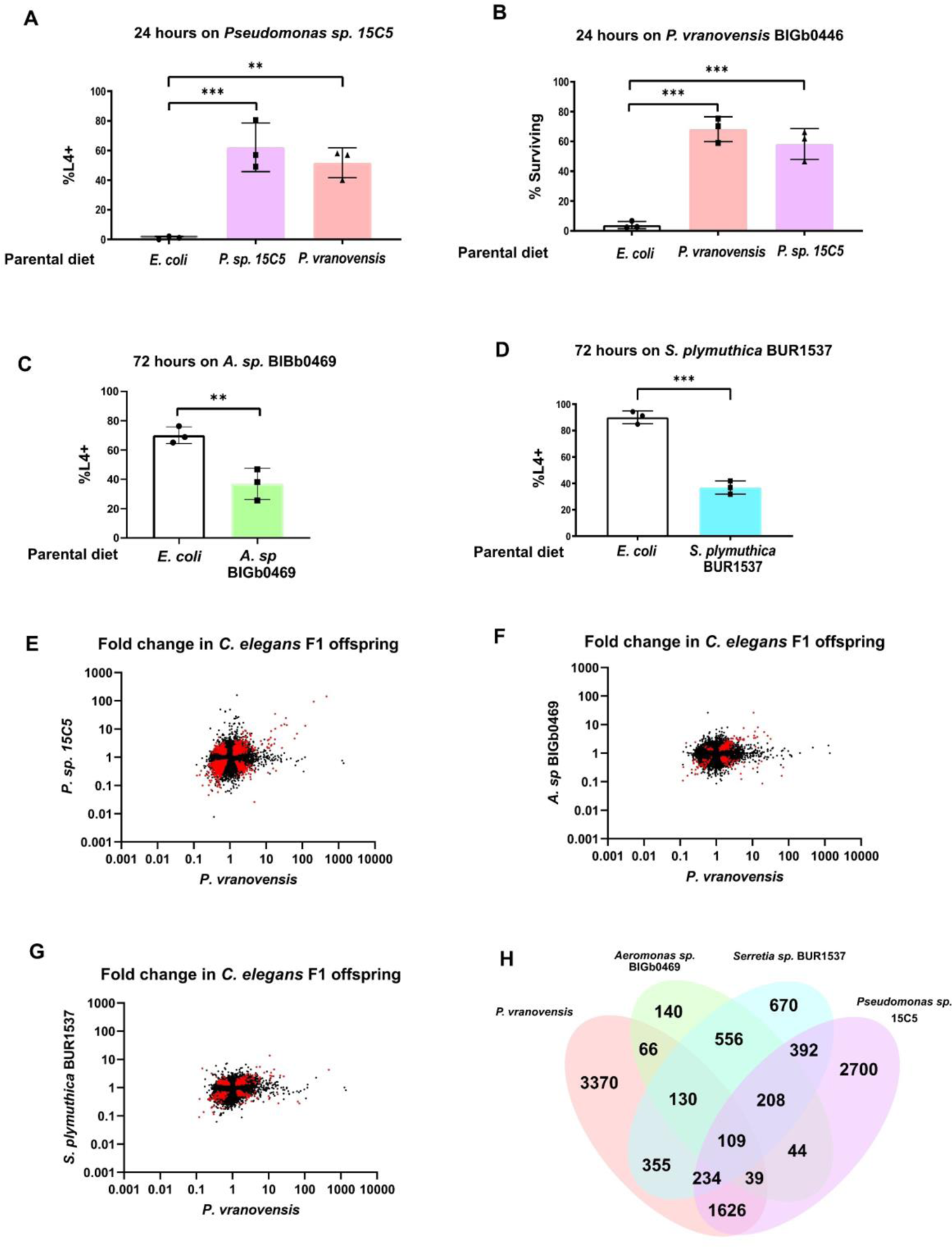
Many of the intergenerational effects of parental exposure to bacterial pathogens on offspring gene expression are pathogen specific. (A) Percent of wild-type *C. elegans* that developed to the L4 larval stage after 48 hours of feeding on *Pseudomonas sp.* 15C5. Data presented as mean values ± s.d. *n* = 3 experiments of >100 animals. (B) Percent of wild-type *C. elegans* surviving after 24 hours of exposure to *P. vranovensis* BIGb0446. Data presented as mean values ± s.d. *n* = 3 experiments of >100 animals. (C) Percent of wild-type *C. elegans* that developed to the L4 larval stage after 48 hours of feeding on *Aeromonas sp.* BIGb0469. Data presented as mean values ± s.d. *n* = 3 experiments of >100 animals. (D) Percent of wild-type *C. elegans* that developed to the L4 larval stage after 48 hours of feeding on *Serratia plymuthica* BUR1537. Data presented as mean values ± s.d. *n* = 3 experiments of >100 animals. (E) Average fold change of genes in F1 progeny of *C. elegans* fed either *Pseudomonas sp.* 15C5 or *P. vranovensis* BIGb0446 when compared to parents fed *E. coli* HB101. Average fold change from three replicates. Red dots represent genes that exhibit statistically significant (*padj* < 0.01) changes in the F1 offspring of parents fed both *Pseudomonas sp.* 15C5 and *P. vranovensis* BIGb0446. (F) Average fold change of genes in F1 progeny of *C. elegans* fed either *Aeromonas sp.* BIGb0469 or *P. vranovensis* BIGb0446 when compared to parents fed *E. coli* HB101. Average fold change from three replicates. Red dots represent genes that exhibit statistically significant (*padj* < 0.01) changes in the F1 offspring of parents fed both *Aeromonas sp.* BIGb0469 and *P. vranovensis* BIGb0446. (G) Average fold change of genes in F1 progeny of *C. elegans* fed either *Serratia plymuthica* BUR1537 or *P. vranovensis* BIGb0446 when compared to parents fed *E. coli* HB101. Average fold change from three replicates. Red dots represent genes that exhibit statistically significant (*padj* < 0.01) changes in the F1 offspring of parents fed both *Serratia plymuthica* BUR1537 and *P. vranovensis* BIGb0446. (H) Venn diagram of the number of genes that exhibit overlapping statistically significant (*padj* < 0.01) changes in expression in F1 progeny of *C. elegans* parents fed each different bacterial species. \*\**p* < 0.01, *** *p* < 0.0001,

In addition to these intergenerational adaptive effects, we also identified two bacterial isolates that activate pathogen response pathways, *Serretia plymutica* BUR1537 and *Aeromonas sp.* BIGb0469 (Samuel et al., 2016; Hellberg et al., 2015), that resulted in intergenerational deleterious effects (Figure 4C-D). Parental exposure of animals to these potential bacterial pathogens did not intergenerationally protect animals against *P. vranovensis* (Supplemental Figure 4). We conclude that parental exposure to some species of *Pseudomonas* can protect offspring from other species of *Pseudomonas*, but that these effects are likely specific to a subset of *Pseudomonas* species and not part of a broad response to gram negative bacterial pathogens.

To determine how different parental bacterial infections affect offspring gene expression patterns, we profiled gene expression in the offspring of *C. elegans* parents exposed to each of *P. vranovensis* BIGb0427, *Pseudomonas sp.* 15C5, *Serretia plymuthica* BUR1537, and *Aeromonas sp.* BIGb0469. We found that only 109 genes exhibit differential expression in the offspring of parents exposed to all four potential pathogens (Figure 4E-H). However, we identified 1,626 genes that are specifically differentially expressed in the offspring of parents exposed to *P. vranovensis* and *Pseudomonas sp*. 15C5 but not in the offspring of parents exposed to *S. plymuthica* BUR1537 or *Aeromonas sp.* BIGb0469 (Figure 4H and Supplemental Table 6). We conclude that parental exposure to bacterial pathogens that elicit enhanced offspring resistance to *P. vranovensis* resulted in distinct changes in offspring gene expression that are not observed when parents are exposed to other gram-winegative bacterial pathogens. Collectively, our results suggest that a majority of the intergenerational effects of a parent’s environment on offspring gene expression are both stress and pathogen specific.

## Discussion

Our findings provide some of the first evidence that the mechanisms underlying intergenerational effects of a parent’s environment on offspring are evolutionarily conserved among different species, are stress-specific, and exhibit deleterious tradeoffs in complex environments. These findings provide a base from which we can compare the numerous different reported observations of multigenerational effects in *C. elegans* to similar effects in other species. For example, we identified 279 genes that exhibited intergenerational regulation of expression in response to the specific stresses of *P. vranovensis* infection or osmotic stress in all species studied (Figure 3). We propose that these genes might be particularly tuned for intergenerational regulation and might similarly be involved in intergenerational responses to stress in other species, including species outside the *Caenorhabditis* genus.

Notably, we found that the expression of these 279 genes in the offspring of parents exposed to either *P. vranovensis* infection or osmotic stress were still differentially expressed in *C. tropicalis* even though parental exposure to these stresses did not appear to affect offspring stress resistance in either assay (Figures 1 and 2). We hypothesize that the molecular consequences of parental stress on offspring, such as changes in the expression of stress response genes, might be more easily identifiable than the specific physiological consequences of parental stress on offspring. In this case we might not have detected the unique phenotypic effects of parental exposure to stress on offspring in *C. tropicalis* using our assay conditions, but such effects might still exist in this species and be related to those observed in other species. Future studies of the phenotypic effects of parental stress on offspring across species will likely shed significant light on how similar molecular mechanisms can mediate different intergenerational responses to stress across evolution.

Consistent with the hypothesis that parental exposure to the same stress might elicit distinct phenotypic effects on offspring in different species via evolutionarily related mechanisms, we found that parental exposure of *C. briggsae* to *P. vranovensis* had a strong deleterious effect on offspring pathogen resistance even though parental exposure of *C. elegans* and *C. kamaaina* to *P. vranovensis* resulted in increased offspring resistance to *P. vranovensis* (Figure 1B). This inversion of an intergenerational effect from a presumed adaptive effect to a presumed deleterious effect correlated with an inversion in the expression of specific pathogen response genes that were previously reported to be required for animals to intergenerationally adapt to *P. vranovensis*, such as *rhy-1* which exhibits increased expression in *C. elegans* and *C. kamaaina* offspring from infected parents but decreased expression in *C. briggsae* offspring from infected parents (Figure 2E). To our knowledge, these findings are the first to suggest that the molecular mechanisms underlying presumed adaptive and deleterious intergenerational effects in different species are evolutionarily related at the gene expression level. These findings suggest that similar observations of presumed intergenerational deleterious effects in diverse species, such as fetal programming in humans, might also be molecularly related to intergenerational adaptive effects in other species. Alternatively, our findings suggest that presumed intergenerational deleterious effects might in fact represent deleterious tradeoffs that are adaptive in other contexts. We expect that a more complete consideration of the evolution of intergenerational effects and the potential relationship between adaptive and deleterious effects will play an important role in understanding how intergenerational effects contribute to organismal resilience in changing environments, what role such effects play in evolution, and how such effects contribute to multiple human pathologies associated with a parent’s environment (Langley-Evans, 2006).

Lastly, the extent to which intergenerational and transgenerational responses to environmental stress represent related, independent, or even mutually exclusive phenomena represents a major outstanding question in the field of multigenerational effects. Evolutionary modelling of intergenerational and transgenerational effects has suggested that different evolutionary pressures favor the evolution of either intergenerational or transgenerational responses under different conditions. Specifically, it has been suggested that intergenerational effects are favored when offspring environmental conditions are predictable from the parental environment (Uller, 2008). Furthermore, it has been speculated that intergenerational adaptations to stress will have costs (Uller, 2008). These costs, such as the costs we observed for animals intergenerational adaptation to osmotic stress (Figure 3), are likely to strongly favor the loss or active erasure of intergenerational effects if the parental environment improves to avoid potential deleterious effects when a stress is no longer present. By contrast, transgenerational effects were found to predominantly be favored when parental environmental cues are unreliable and the maintenance of information across many generations might be worth the potential costs (Uller et al., 2015). Our findings support a model in which intergenerational and transgenerational effects represent potentially distinct phenomena. Specifically, multiple of the intergenerational responses to different stresses studied here were previously reported to be intergenerational in nature and only last for a single generation (Burton et al., 2020, 2017; Hibshman et al., 2016; Willis et al., 2021). Our studies provide further evidence that the effects of the parental exposure to osmotic stress and *P. vranovensis* infection are predominantly intergenerational in nature as we did not detect any conserved transgenerational changes in gene expression in response to either stress (Figure 2). We strongly suspect that future studies into the mechanisms regulating these intergenerational effects will shed significant light on how intergenerational effects on gene expression are lost and/or erased. In addition, we expect that similar studies of transgenerational effects will potentially elucidate the circumstances under which animals decide environmental information might be worth maintaining despite any potential tradeoffs and if the growing number of transgenerational effects observed in *C. elegans* are similarly evolutionarily conserved.

Lastly, future studies of intergenerational effects will be critical in determining the extent to which the mechanisms that mediate intergenerational effects are conserved outside of *Caenorhabditis* and if similar mechanisms to those uncovered in *C. elegans* mediate the numerous different adaptive and deleterious intergenerational effects that have been reported in diverse taxa ranging from the intergenerational development of wings in aphids (Vellichirammal et al., 2017) to fetal programming and the role it plays in disease in humans (Langley-Evans, 2006).

## Methods

### Strains

*C. elegans* strains were cultured and maintained at 20 °C unless noted otherwise. The Bristol strain N2 was the wild-type strain. Wild isolate strains used in the main figures of this study: N2 (*C. elegans*), AF16 (*C. briggsae*), JU1373 (*C. tropicalis*), and QG122 (*C. kamaaina*). Wild-isolate strains used in supplemental figures of this study: MY1 (*C. elegans*), PS2025 (*C. elegans*), CX11262 (*C. elegans*), JU440 (*C. elegans*), JU778 (*C. elegans*), JU1213 (*C. elegans*), LKC34 (*C. elegans*), JU1491 (*C. elegans*), EG4724 (*C. elegans*), KR314 (*C. elegans*), SX1125 (*C. briggsae*), and JU1348 (*C. briggsae*). Mutant alleles used in this study: *osm-8(n1518)* and *Cbr-gpdh-2(syb2973)*.

### P. vranovensis *survival assays.*

*P. vranovensis* BIGb0446 or *Pseudomonas sp.* 15C5 was cultured in LB at 37 °C overnight. 1 mL of overnight culture was seeded onto 50 mm NGM agar plates and dried in a laminar flow hood (bacterial lawns completely covered the plate such that animals could not avoid the pathogen). All plates seeded with BIGb0446 or 15C5 were used the same day they were seeded. Young adult animals were placed onto 50 mm NGM agar plates seeded with 1 mL either *E. coli* HB101, *P. vranovensis* BIGb446, or *Pseudomonas sp.* 15C5 for 24 h at room temperature (22 °C). Embryos from these animals were collected by bleaching and placed onto fresh NGM agar plates seeded with BIGb0446. Percent surviving were counted after 24 h at room temperature (22 °C) unless otherwise noted.

### Osmotic stress and P. vranovensis *multiple stress adaptation assays*

Young adult animals that were grown on NGM agar plates seeded with *E. coli* HB101 were collected and transferred to new 50 mM NaCl control plates seeded with *E. coli* HB101, 300 mM NaCl plates seeded with *E. coli* HB101, 50 mM NaCl control plates seeded with *P. vranovensis* BIGb0446, or, 300 mM NaCl plates seeded with *P. vranovensis* BIGb0446. Animals were grown for 24 hours at room temperature (22 °C). Embryos from these animals were collected by bleaching and transferred to new 500 mM NaCl plates seeded with *E. coli* HB101 or 50 mM NaCl plates seeded with *P. vranovensis* BIGb0446. Percent of animals developing or surviving was scored after 24 hours at room temperature as previously described in Burton et al., 2017 and Burton et al., 2020.

### Preparation of N. parisii spores

Spores were prepared as described previously (Willis et al., 2021). In brief, large populations of *C. elegans* N2 were infected with microsporidia spores. Infected worms were harvested and mechanically disrupted using 1 mm diameter Zirconia beads (BioSpec). Resulting lysate was filtered through 5 μm filters (Millipore Sigma**™**) to remove nematode debris. Spore preparations were tested for contamination and those free of contaminating bacteria were stored at −80°C.

### N. parisii infection assays and multiple stress adaptation assays

P0 populations of 2500 animals were mixed with 1 ml of 10X saturated *E. coli* OP50-1 or *P. vranovensis* and a low dose of *N. parisii* spores (see Table 1) and plated on a 10 cm plate. This low dose limited the detrimental effects on animal fertility that are observed with higher doses, while ensuring most animals were still infected. F1 populations of 1000 animals were mixed with 400 μl of 10X saturated *E. coli* OP50-1 and a high dose of *N. parisii* spores (see Table 1) and plated on a 6 cm plate.

**Table 1.** *Details of* N. parisii *doses* employed.

| *N. parisii*<br>dose | Plate concentration<br>(spores/cm<sup>2</sup>) | Millions of spores used |  |
| --- | --- | --- | --- |
|  |  | 6 cm plate | 10 cm plate |
| Low | ~32,000 |  | 2.5 |
| High | ~88,000 | 2.5 |  |

To test for inherited immunity to *N. parisii* in *C. elegans, C. briggsae*, *C. tropicalis* and *C. kamaaina*, synchronized animals were infected from the L1 larval stage with a low dose of *N. parisii*. *C. elegans and C. briggsae* were grown for 72 hours at 21°C; *C. tropicalis* and *C. kamaaina* were grown for 96 hours at 21°C. Ten percent of total P0 animals were fixed in acetone for DY96 staining, as described below. Embryos from the remaining animals were collected by bleaching and synchronized by hatching overnight in M9. Resulting F1 animals were infected from the L1 larval stage with a high dose of *N. parisii. C. elegans* and *C. briggsae* were fixed at 72 hours post infection (hpi) at 21°C; *C. tropicalis* and *C. kamaaina* were fixed at 96 hpi at 21°C.

For multiple stress adaptation assays using *N. parisii* and osmotic stress, animals were grown on NGM agar plates seeded with 10X saturated *E. coli* OP50-1 until the L4 stage. Next, animals were collected and mixed with 1 ml of either *E. coli* OP50-1 alone or supplemented with a low dose of *N. parisii* spores and plated on either 50 mM NaCl or 250 mM NaCl plates. Animals were grown for 24 hours at 21°C. Embryos from these animals were collected by bleaching. To test adaptation to osmotic stress, 2000 F1 embryos were transferred to 420 mM NaCl plates seeded with *E. coli* OP50-1. Percentage of animals hatched was scored after 48 hours at 21°C, as previously described in Burton et al., 2017 and Burton et al., 2020. To test adaptation to *N. parisii*, the remaining embryos were synchronized by hatching overnight in M9. Resulting F1 animals were either not infected as controls, or infected at the L1 larval stage with a high dose of *N. parisii*. Animals were fixed after 72 hours at 21°C for DY96 staining and analysis.

For multiple stress adaptation assays using *N. parisii* and *P. vranovensis*, animals were grown on NGM agar plates seeded with *E. coli* OP50-1 until the L4/young adult stage. Next, animals were collected and mixed with 1 ml of either *E. coli* OP50-1 alone or *E. coli* OP50-1 supplemented with a low dose of *N. parisii* spores, or 1 ml of *P. vranovensis* BIGb0446 alone or *P. vranovensis* BIGb0446 supplemented with a low dose of *N. parisii* spores. Animals were plated on NGM and grown for 24 hours at 21°C. Embryos from these animals were collected by bleaching. To test adaptation to *P. vranovensis*, 2000 F1 embryos were transferred to new NGM plates seeded with *P. vranovensis* BIGb0446. Percentage of animals surviving was scored after 24 hours at 21°C as previously described in Burton et al., 2017 and Burton et al., 2020. To test adaptation to *N. parisii*, the remaining embryos were synchronized by hatching overnight in M9. Resulting F1 animals were either not infected as controls, or infected from the earliest larval stage with a high dose of *N. parisii*. Animals were fixed after 72 hours at 21°C for DY96 staining and analysis.

### Fixation and staining of *N. parisii* infection

Worms were washed off plates with M9 and fixed in 1 ml acetone for 10 min at room temperature, or overnight at 4°C. Fixed animals were washed twice in 1 ml PBST (phosphate buffered saline (PBS) containing 0.1% Tween-20) before staining. Microsporidia spores were visualized with the chitin-binding dye Direct Yellow (DY96). For DY96 staining alone, animals were resuspended in 500 μl staining solution (PBST, 0.1% sodium dodecyl sulfate [SDS], 20 ug/ml DY96), and rotated at 21°C for 30 min in the dark. DY96-stained worms were resuspended in 20 μl EverBrite™ Mounting Medium (Biotium) and mounted on slides for imaging. Note: to pellet worms during fixation and staining protocols, animals were centrifuged for 30 seconds at 10,000 xg.

### Image analysis of N. parisii infection

Worms were imaged with an Axioimager 2 (Zeiss). DY96-stained worms were imaged to determine number of embryos per worm. Worms possessing any quantity of intracellular DY96-stained microsporidia were considered infected. Precise microsporidia burdens were determined using ImageJ/FIJI (Schindelin et al., 2012). For this, each worm was defined as an individual ‘region of interest’ and fluorescence from GFP (DY96-stained microsporidia) subject to ‘threshold’ and ‘measure area percentage’ functions on ImageJ. Images were thresholded to capture the brighter signal from microsporidia spores, while eliminating the dimmer GFP signal from worm embryos. Final values are given as % fluorescence for single animals.

### Preparation of OP50 for plating worms

One colony of *E. coli* strain OP50 was added to 100mL of LB and grown overnight at room temperature then stored at 4°C. 1 or 5 drops of HB101 were added to 6 or 10 cm plates of NGM, respectively, to use for growing worm strains and recovering them from starvation.

### Preparation of HB101 for liquid culture

One colony of *E. coli* strain HB101 was added to a 5mL starter culture of LB with streptomycin and grown for 24 hours at 37°C. The starter cultures were then added to a 1L culture of TB and grown for another 24 hours at 37°C. The bacteria was centrifuged for 10 min at 5000 RPM to form a pellet. After being weighed, the bacteria was then resuspended in S-complete to create a 10x (250mg/mL) stock that was stored at 4°C. Further dilutions with S-complete were used to create the dilutions for each condition in this experiment.

### Dietary restriction/dilution series cultures

For *C. elegans, C. briggsae* and *C. tropicalis*, 10 L4 hermaphrodite worms were picked onto 3 10 cm plates seeded with OP50, and for *C. kamaaina* 10 L4 females and ∼20 males were picked onto 3 10 cm plates. For all species, adults were removed after 24 hours. *C. elegans* and *C. briggsae* were grown for 96 hours before bleaching and *C. tropicalis* and *C. kamaaina* were grown for 120 hours before bleaching due to slower growth and longer generation time. After bleaching, worms were aliquoted into 100 mL cultures of S-complete at 1 worm/100 μL with a concentration of 25mg/mL, 12.5mg/mL 6.25mg/mL, 3.13mg/mL or 1.6 mg/mL of HB101 and kept in 500mL flasks in shaking incubators at 20°C and 180 RPM. Worms were grown in these cultures for 96 hrs (*C. elegans*), 102 hrs (*C. briggsae*) or 120 hrs (*C. tropicalis* and *C. kamaaina*) before being bleached and prepared for starvation cultures. Due to slow development and inability to properly scale up in liquid culture, 1.6mg/mL cultures for *C. briggsae* and 1.6 and 3.13 mg/mL cultures for *C. kamaaina* were excluded from the rest of this experiment.

### Starvation cultures

After bleach, embryos were placed into 5mL virgin S-basal cultures in 16 mm glass test tubes on a roller drum at 20°C at 1 worm/μL. Worms were aliquoted out of this culture using micropipettes for further assays.

### Measuring L1 size

24 hours after bleach (∼12 hours after hatch), 1000 L1s were pipetted out of the starvation cultures, spun down in 15mL plastic conical tubes by centrifuge for 1 min at 3000 RPM then plated onto unseeded 10cm NGM plates. L1s were imaged with a Zeiss Discovery.V20 stereomicroscope at 77x and measured using Wormsizer (Moore et al., 2013). Ad libitum concentration was defined as 25mg/mL and dietary restriction concentration was determined based on what concentration of HB101 produced the largest average L1 size for each strain. For *C. elegans*, this was 3.13mg/mL, and 8-fold dilution from ad libitum and consistent with previous determinations for dietary restriction in *C. elegans* (Hibshman et al., 2016). For *C. briggsae*, peak L1 size varied between 12.5mg/mL and 6.25 mg/mL depending on replicate. We chose to use 6.25 mg/mL as the dietary restriction concentration to be consistent with replicates that were already being processed. The peak L1 size and determination of dietary restriction for *C. tropicalis* was 6.13 mg/mL*. C. kamaaina* did not show a significant change in L1 size across conditions and was ultimately excluded from the brood size assay due to difficulty interpreting effects of starvation on brood size in a male-female strain.

### L1 size statistics

A linear mixed effects model was performed on the L1 size data to see if there was a significant effect of HB101 concentration on L1 size. The lme4 package in R studio was used to perform this linear mixed effects test. The function lmer() was used on data from each species, for example: • lmer(length ∼ condition + (1 | replicate) + (1 | replicate:condition), data = C_elegans), “length” is the length in microns of each individual worm, “condition” is the fixed effect of the concentration of HB101, “1 | replicate” is the addition of the random effect of replicate to the model, “1 | replicate:condition” is the addition of the random effect per combination of replicate and condition, and “data” is the primary spreadsheet restricted by the species of interest.

### Gene orthology inference among species

To identify one-to-one orthologues across the four species, we downloaded protein and GFF3 files for *C. elegans*, *C. briggsae*, and *C. tropicalis* genomes from WormBase (Harris et al., 2020) (version WS275) and for the *C. kamaaina* genome from caenorhabditis.org (version v1). We assessed gene set completeness using BUSCO (Simão et al., 2015) (version 4.0.6; using the parameter *-m proteins*) using the ‘nematoda_odb10’ lineage dataset. For each species, we selected the longest isoform for each protein-coding gene using the agat_sp_keep_longest_isoform.pl script from AGAT (Jacques Dainat et al., 2021) (version 0.4.0). Filtered protein files were clustered into orthologous groups (OGs) using OrthoFinder (Emms and Kelly, 2019) (version 2.4.0; using the parameter *-og*) and one-to-one OGs were selected.

### F1 and F3 sample collection for RNA-seq

Young adult animals grown on NGM agar plates seeded with *E. coli* HB101 were collected and transferred to new plates seeded with either control plates (50 mM NaCl) seeded with *E. coli* HB101, *P. vranovensis* BIGb0446, *P. vranovensis* BIGb0427, *S. plymuthica* BUR1537, *Pseudomonas sp.* 15C5, *Aeromonas sp.* BIGb0469, or plates containing 300 mM NaCl seeded with *E. coli* HB101. Animals were grown for 24 hours at room temperature (22 °C). Embryos from these animals were collected by bleaching and immediately frozen in 1 mL Trizol.

### Analysis of RNA-Seq data

RNA libraries were prepared and sequenced by BGI TECH SOLUTIONS using 100PE DNBseq Eukaryotic Transcriptome service. Quality controlled and adapter trimming of RNA reads were performed using fastp-v4.20.0 (Chen et al., 2018) (--qualified_quality_phred 20 --unqualified_percent_limit 40 --length_required 50 -- low_complexity_filter --complexity_threshold 30 --detect_adapter_for_pe --correction -- trim_poly_g --trim_poly_x \ --trim_front1 2 --trim_tail1 2 --trim_front2 2 --trim_tail2 2) **1)**.

Next, reads were aligned using STAR-2.7.1a (Dobin et al., 2013) (--alignSJoverhangMin 8 -- alignSJDBoverhangMin 1 --outFilterMismatchNmax 999 -- outFilterMismatchNoverReadLmax 0.04 --alignIntronMin 10 --alignIntronMax 1000000 -- alignMatesGapMax 1000000 --outFilterType BySJout --outFilterMultimapNmax 10000 -- winAnchorMultimapNmax 50 --outMultimapperOrder Random) **2)** against the genome of *C. elegans* WS275, *C. briggsae* WS275, *C. tropicalis* WS275 and the *C. kamaaina* genome obtained from caenorhabditis.org. Read counts were obtained using subread-2.0.0 (-M -O -p --fraction -F GTF -a -t exon -g gene_id) (Liao et al., 2014) **3)** using the annotation for *C. elegans* PRJNA13758.WS275, *C. briggsae* PRJNA10731.WS275, *C. tropicalis* PRJNA53597.WS275, and *C. kamaaina* Caenorhabditis_kamaaina_QG2077_v1. Counts were imported into R and differential gene expression analysis was performed with DESeq2 (FDR <0.01) (Love et al., 2014).

For comparisons made between different species, genes were subsetted to include only those 7,587 single copy ortholog groups that were identified between the four species. In addition to the 7,203 genes that were identified as single copy ortholog groups by OrthoFinder, the 7,587 contain an additional 385 ortholog groups that were identified as having more than one ortholog in one out four of the species but where all but one of the multiple orthologs had no observable expression in any of the samples collected.

For the comparison between the stress response and gene expression during embryo development, data was downloaded from Boeck et al., 2016 and imported in R with raw counts from this study. The range of embryo expression for each gene was considered as one standard deviation plus / minus the mean of regularised log normalised counts across all embryo time points. DEGs from the stress experiments where the regularised log normalised counts for one or both of the comparison samples (for all replicates) were outside of the embryo range were considered unlikely to be caused by developmental timing.

### L4+ developmental rate assays

Young adult animals that were grown on NGM agar plates seeded with *E. coli* HB101 were collected and transferred to new plates seeded with *E. coli* HB101, *Pseudomonas sp*. 15C5, *S. plymuthica* BUR1537, or *Aeromonas sp*. BIGb0469. Animals were grown for 24 hours at room temperature (22°C). Embryos from these animals were collected by bleaching and transferred to new plates seeded with 1 mL of *E. coli* HB101 *Pseudomonas sp*. 15C5, *S. plymuthica* BUR1537, or *Aeromonas sp*. BIGb0469. Percent of animals that reached the L4 larval stage was scored after either 48 hours or 72 hours at 22°C.

### Identification of Pseudomonas sp. 15C5 and S. plymuthica BUR1537

Samples of rotting fruit and vegetation were collected from around Cambridge (UK) in 50 mL vials. For isolation of wild bacteria, the samples were homogenized and resuspended in M9 and plated on LB Agar, Nutrient Agar, or Actinomycete Isolation Agar plates and grown at either 37°C or 30°C for 24 hours. Single colonies were isolated from the plates and grown in LB or Nutrient Broth at the same temperature overnight. Stocks were frozen and stored at −80 °C in 20% glycerol. 1,537 total isolates were obtained and frozen. *C. elegans* embryos were placed onto NGM agar plates seeded with each of the 1,537 bacterial isolates. Bacterial isolates that caused substantial delays in animal development or lethality were further analyzed for isolates where parental exposure to the isolate for 24 hours modified offspring phenotype when compared to offspring from parents fed the normal laboratory diet of *E. coli* HB101. Bacterial genus and species were identified by 16S rRNA profiling and sequencing.

### RNAi in C. kamaaina

dsDNA corresponding to the *C. kamaaina* orthologues of *cysl-1*, *rhy-1*, *mek-2* and *gpdh-2* was synthesized and cloned into the L4440 vector by GENEWIZ (Takeley, UK). Vectors were transformed in *E. coli* HT115. *C. kamaaina* embryos were collected by bleaching and placed onto NGM agar plates containing 1 mM IPTG that were seeded with E. coli HT115 transformed with either the L4440 empty vector or each of the new vectors and grown at room temperature (22°C) for 48 hours. After 48 hours animals were transferred to new 50 mM NaCl control plates seeded with *E. coli* HB101, 300 mM NaCl plates seeded with *E. coli* HB101, or 50 mM NaCl control plates seeded with *P. vranovensis* BIGb0446. Animals were grown for 24 hours at room temperature (22 °C). Embryos from these animals were collected by bleaching and transferred to new 500 mM NaCl plates seeded with *E. coli* HB101 or 50 mM NaCl plates seeded with *P. vranovensis* BIGb0446. Percent of animals developing or surviving was scored after 24 hours at room temperature as previously described in Burton et al., 2017 and Burton et al., 2020.

### Statistics and reproducibility

Sample sizes for experiments involving *C. elegans* were selected based on similar studies from the literature and all animals from each genotype and condition were selected and analyzed randomly. All replicate numbers listed in figure legends represent biological replicates of independent animals cultured separately, collected separately, and analyzed separately. Unpaired two-tailed Student’s *t*-test was used for Fig. 1B, 1D, 1F, 2E, 2J, 4C, 4D, and Supplemental Fig. 2. Two-way ANOVA was used for Fig. 1C, 1E, 3A-G, and Supplemental Fig. 1. One-way ANOVA was used for Fig. 4A, 4B, and Supplemental Fig. 3. ^*^= *P* < 0.05, ^**^ = *P* < 0.01, ^***^ = *P* < 0.001, ^****^*P* < 0.0001. The experiments were not randomized. The investigators were not blinded to allocation during experiments and outcome assessment. Source data for all graphs can be found in the Source Data Supplemental Table.

### Data availability

RNA-seq data that support the findings of this study have been deposited at NCBI GEO and are available under the accession code GSE173987.

## Supporting information

Statistics Source Data

## Acknowledgments

We would like to thank Buck Samuel and Marie-Anne Felix for bacterial isolates. We also thank Matt Rockman and Luke Noble for pre-publication access to the genome of *C. kamaaina*. We would also like to thank Marie-Anne Felix and and the *Caenorhabditis* Genetic Center, which is funded by the NIH National Center for Research Resources (NCRR), for *Caenorhabditis* strains. N.O.B. is funded by a Next Generation Fellowship from the Centre for Trophoblast Research. K.F. and L.R.B. were funded by the National Institutes of Health (GM117408, L.R.B). A.W. and A.R. were funded by the Natural Sciences and Engineering Research Council of Canada (Grant #522691522691) and an Alfred P. Sloan Research Fellowship FG2019-12040 (to A.W.R.). This work was also supported by Cancer Research UK (C13474/A18583, C6946/A14492) and the Wellcome Trust (104640/Z/14/Z, 092096/Z/10/Z) grants to E.A.M.

## Author Contributions

N.O.B., A.W., A.R., K.F., and L.R.B. designed the experiments. N.O.B, A.W., J.P., F.B., L.S., K.F., A.R., L.R.B, and E.A.M analysed the data. N.O.B., A.W., K.F., and F.B. performed the experiments. N.O.B. conceived the project and wrote the manuscript.

---

**Supplemental Figure 1.**
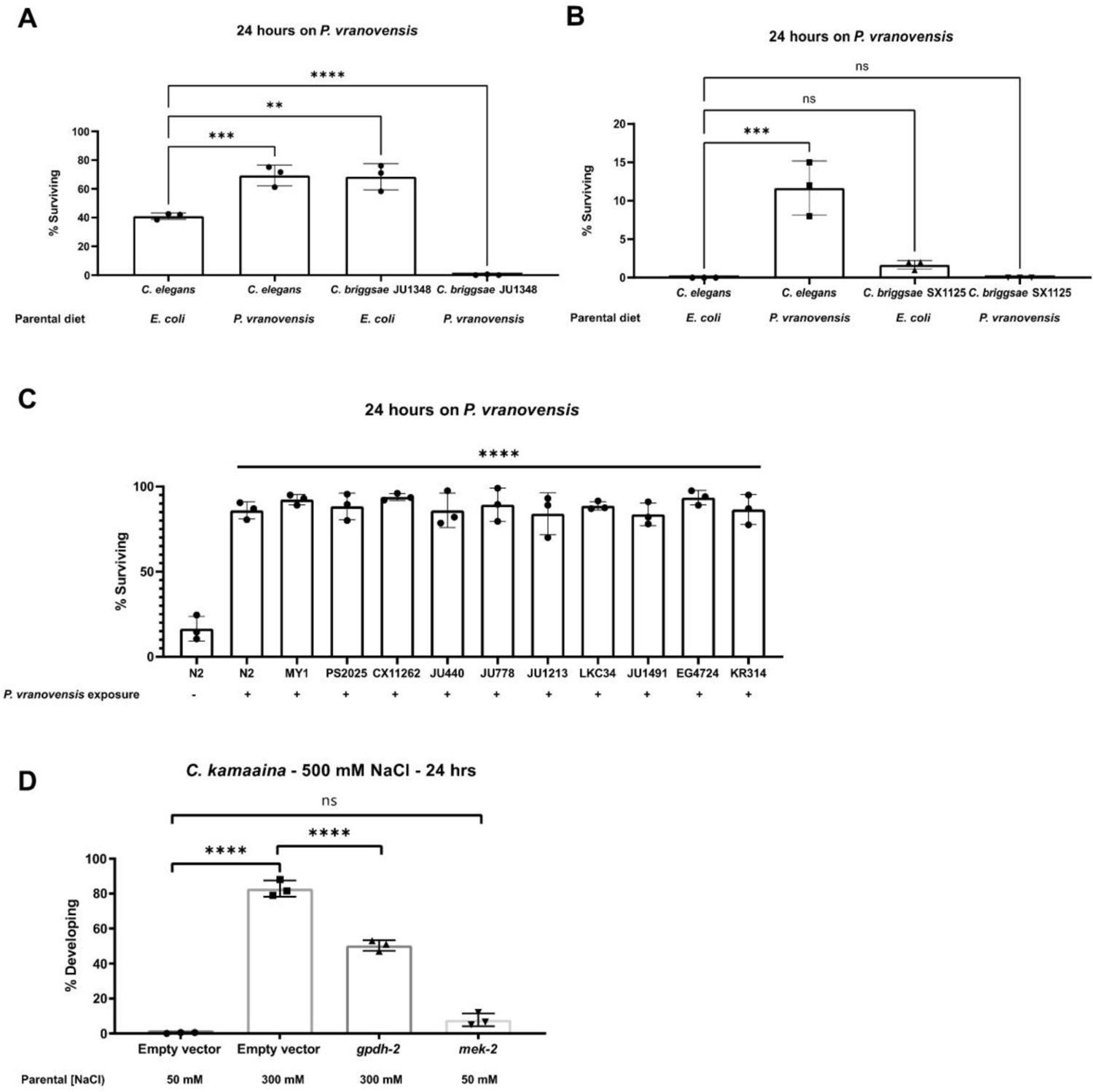
Intergenerational responses to environmental stress are conserved in wild isolates of *Caenorhabditis* species. (a) Percent of wild-type *C. elegans* (N2) and *C. briggsae* (JU1348) animals surviving after 24 hours on plates seeded with *P. vranovensis* BIGb0446. Data presented as mean values ± s.d. *n* = 3 experiments of >100 animals. (b) Percent of wild-type *C. elegans* (N2) and *C. briggsae* (SX1125) animals surviving after 24 hours on plates seeded with *P. vranovensis* BIGb0446. Data presented as mean values ± s.d. *n* = 3 experiments of >100 animals. (c) Percent of wild-type *C. elegans* isolates surviving after 24 hours on plates seeded with *P. vranovensis* BIGb0446. Data presented as mean values ± s.d. *n* = 3 experiments of >100 animals. (d) Percent of wild-type *C. kamaaina* animals mobile and developing at 500 mM NaCl after 24 hours. Data presented as mean values ± s.d. *n* = 3 experiments of >100 animals.

**Supplemental Figure 2.**
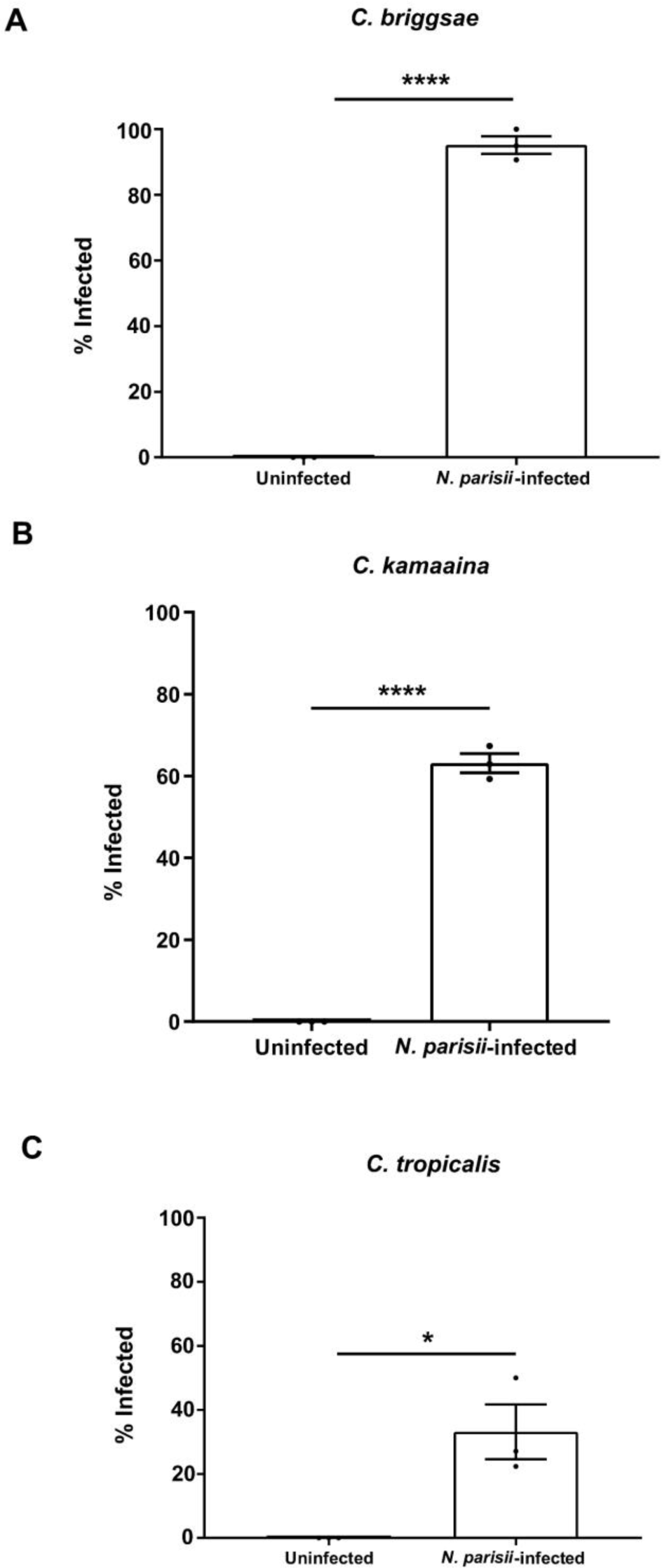
*N. parisii* infects *C. briggsae, C. kamaaina, and C. tropicalis*. *N. parisii* infects *C. briggsae, C. kamaaina, and C. tropicalis*. Percent of animals exhibiting detectable infection by *N. parisii* as determined by DY96 staining after 72 h for *C. elegans* and *C. briggsae*, or 96 h for *C. kamaaina* and *C. tropicalis*. Data presented as mean values ± s.e.m. *n* = 3 experiments of 68-115 animals (1A), 27-102 animals (1B), 38-104 animals (1C).

**Supplemental Figure 3.**
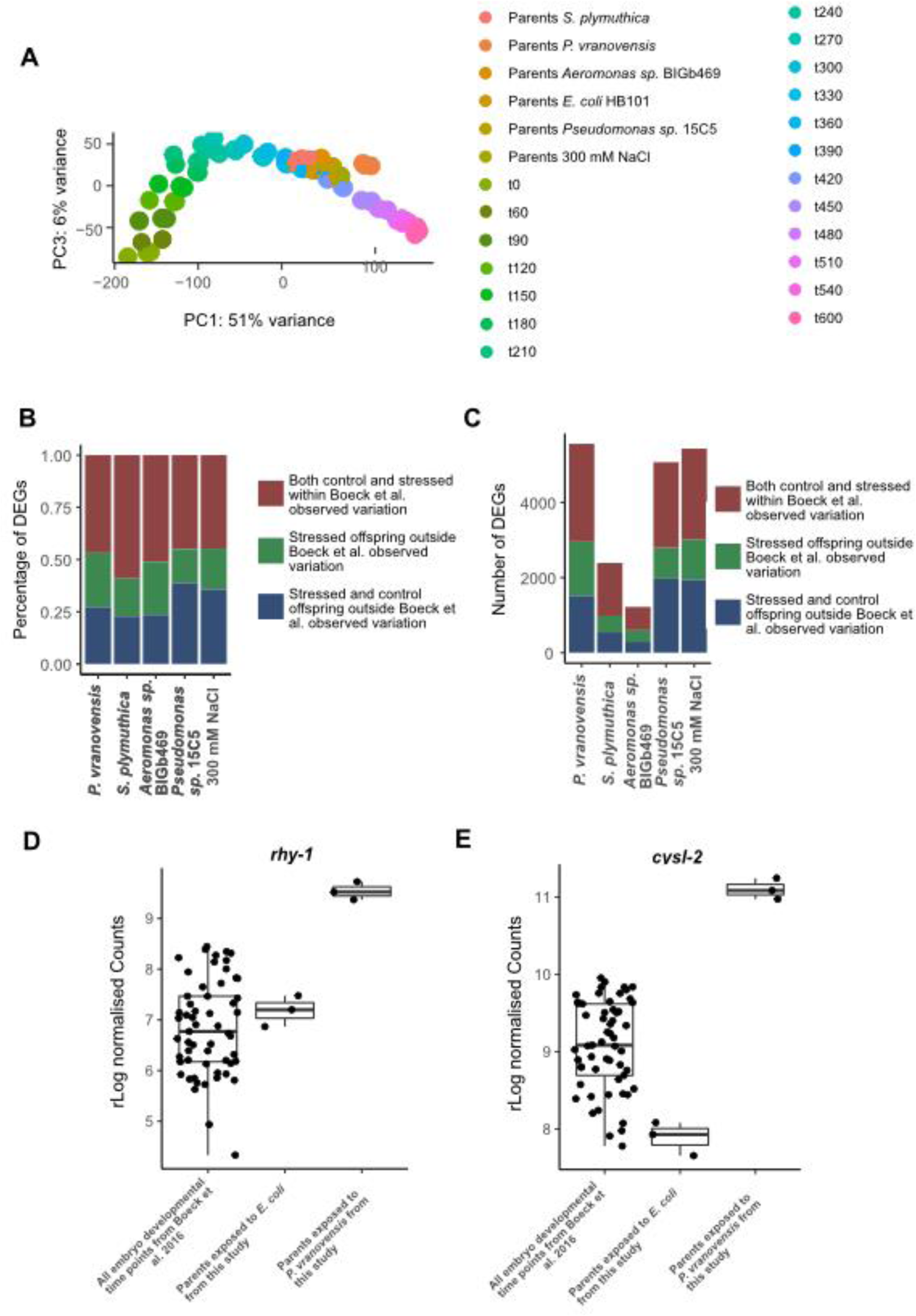
Differences in developmental timing are insufficient to explain a majority of the observed differences in gene expression in the offspring of stressed parents. (A) PCA of gene expression from Boeck et al. (2016) compared to RNA-seq data reported in the study. Time points of development are in minutes, t60 = 60 minutes post fertilization. (B) Percentage of genes differentially expressed in the offspring of parents exposed to different stresses that exhibit DESeq2 normalized counts that fall within or outside one standard deviation of the average normalized counts observed throughout all developmental timepoints from Boeck et al. (2016). (C) Total number of genes differentially expressed in the offspring of parents exposed to different stresses that exhibit DESeq2 normalized counts that fall within or outside one standard deviation of the average normalized counts observed throughout all developmental timepoints from Boeck et al. (2016). (D) *rhy-1* normalized counts from all time points during development from Boeck et al. (2016), the offspring of parents exposed to *E. coli* HB101 (this study), or the offspring of parents exposed to *P. vranovensis* (this study). (E) *cysl-2* normalized counts from all time points during development from Boeck et al. (2016), the offspring of parents exposed to *E. coli* HB101 (this study), or the offspring of parents exposed to *P. vranovensis* (this study).

**Supplemental Figure 4.**
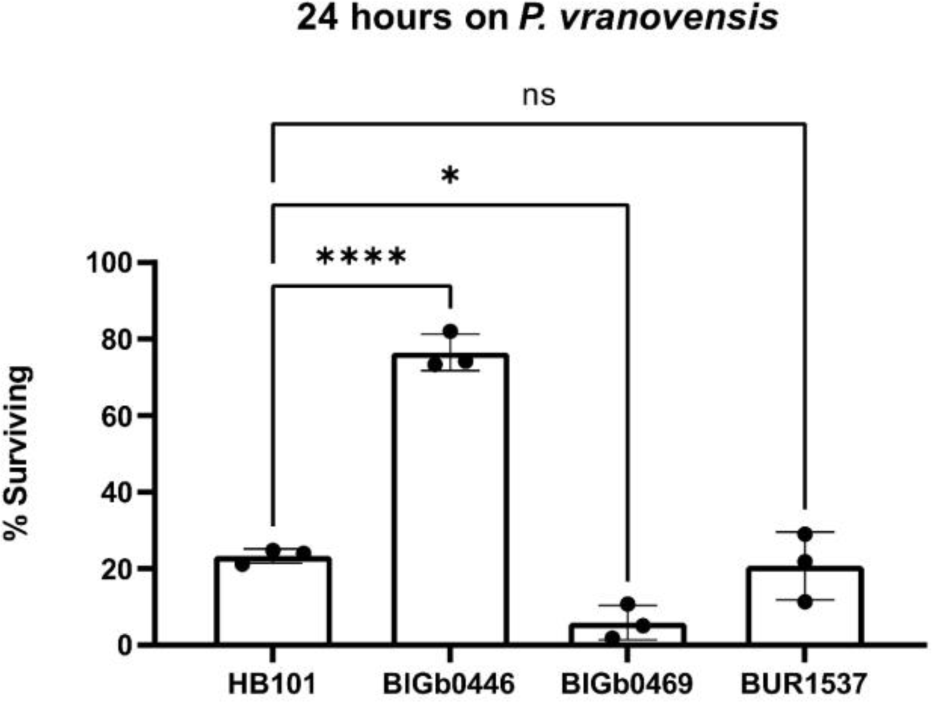
Parental exposure to *Aeromonas sp.* BIGb0469 and *S. plymuthica* BUR1537 does not protect offspring from *P. vranovensis.* Percent of wild-type *C. elegans* (N2) animals surviving after 24 hours on plates seeded with *P. vranovensis* BIGb0446. Data presented as mean values ± s.d. *n* = 3 experiments of >100 animals.

